# Microbiome of psyllids of the family Aphalaridae, including *Aphalara itadori*, a potential biocontrol agent against *Reynoutria* spp

**DOI:** 10.1101/2024.03.28.587303

**Authors:** Kyosuke Nishino, Hiromitsu Inoue, Yuu Hirose, Atsushi Nakabachi

## Abstract

Several European and North American countries have started releasing the Japanese knotweed psyllid *Aphalara itadori* (Hemiptera: Aphalaridae) to control the Japanese knotweed *Reynoutria japonica* (Caryophyllales: Polygonaceae) and its relatives, which are among the worst invasive exotic plants. However, establishing populations of the current Kyushu and Hokkaido strains in the field has not been successful, desiring new lineages. Moreover, little is known about the microbiome of the current strains, which potentially impacts properties as biocontrol agents. Hence, this study analyzed the microbiota of an *A. itadori* strain newly collected on Honshu Island, along with related species of the family Aphalaridae, using amplicon sequencing of 16S rRNA genes. The *A. itadori* symbionts were further located using fluorescence *in situ* hybridization. The results demonstrated that the analyzed *A. itadori* strain has a dual symbiotic system with “*Candidatus* Carsonella ruddii” (Gammaproteobacteria: Oceanospirillales) and *Sodalis* sp. (Gammaproteobacteria: Enterobacterales) harbored in the bacteriome, suggesting their evolutionarily stable mutualistic relationships with *A. itadori*. The central area of the bacteriome harboring *Sodalis* appeared to comprise uninucleate bacteriocytes with nuclei larger than those of bacteriocytes for *Carsonella*, rather than a syncytium with smaller nuclei as previously reported for various psyllid lineages. No known plant pathogens or manipulators of insect reproduction were identified in the analyzed strain, indicating its suitability as a biocontrol agent, posing a minimum risk to the ecosystem. Besides distinct *Carsonella* lineages, the analysis identified *Sodalis* independently acquired by *Craspedolepta miyatakeai*, and an ambiguous *Enterobacterales* symbiont in *Epheloscyta kalopanacis*. Only *Carsonella* was identified in *Togepsylla matsumurana*.

## Introduction

Invasive alien species impose a significant impact on native flora and fauna. Among the most devastating examples are the Japanese knotweed *Reynoutria* (= *Fallopia*) *japonica* (Caryophyllales: Polygonaceae), the giant knotweed *Reynoutria sachalinensis*, and their hybrid, the Bohemian knotweed *Reynoutria* × *bohemica*, all of which originated in East Asia (Shaw *et al*., 2009; Djeddour and Shaw, 2010; Andersen *et al*., 2016). They were intentionally introduced from Japan and spread as favored gardening plants in European and North American countries in the 19^th^ century (Djeddour and Shaw, 2010). However, due to the rapid growth potential and tolerance to diverse environmental conditions, they have established dense colonies in numerous habitats, driving out indigenous plants and damaging artificial structures, including buildings, pavements, and waterways, making them now considered among the worst invasive exotic plants in many countries (Shaw *et al*., 2009; Djeddour and Shaw, 2010; Andersen *et al*., 2016). Because they quickly grow back asexually from tiny fragments of rhizomes forming large and deep underground networks, it is difficult to eradicate these knotweeds from invaded areas. As one of the measures to control their populations, several countries, including the United Kingdom, Canada, the United States, and the Netherlands, have started releasing the Japanese knotweed psyllid *Aphalara itadori* (Insecta: Hemiptera: Sternorrhyncha: Psylloidea: Aphalaridae: Aphalarinae), native to Japan, as a promising biological control agent (Shaw *et al*., 2009; Djeddour and Shaw, 2010; Andersen *et al*., 2016; Grevstad *et al*., 2022). Because *A*. *itadori* specifically feeds on *Reynoutria* spp., causing their defoliation and reduction of the growth and root storage, this insect species is anticipated to control invasive knotweeds safely and effectively. The two *A*. *itadori* strains introduced thus far were originally collected on the southernmost and northernmost of Japan’s four main islands, Kyushu and Hokkaido. The Kyushu strain was chosen because the invasive clone of *R. japonica* is believed to have originated in Kyushu (Djeddour and Shaw, 2010). The Hokkaido stain was adopted as this strain is thought to be effective against *R. sachalinensis* (Grevstad *et al*., 2022), which occurs mainly on Hokkaido (Shaw *et al*., 2009). However, establishing populations of these strains in the field has not been as successful as anticipated. Moreover, the hybrid created from the Kyushu and Hokkaido strains aiming to increase genetic and phenotypic variations failed to improve its performance as a biocontrol agent (Fung *et al*., 2020). Therefore, newly established *A*. *itadori* strains distinct from the current Kyushu and Hokkaido strains are desirable (Camargo *et al*., 2022).

Psyllids (Psylloidea), involving approximately 4,000 described species (Burckhardt *et al*., 2021), are insects feeding on phloem sap with their needle-shaped mouthparts (Hodkinson, 1974; Burckhardt *et al*., 2021), as do other insect groups belonging to the suborder Sternorrhyncha (International Aphid Genomics Consortium, 2010; Nakabachi and Miyagishima, 2010; Tamborindeguy *et al*., 2010). Through this feeding habit, some psyllid lineages transmit plant pathogens, including “*Candidatus* Liberibacter spp.” (Alphaproteobacteria: Rhizobiales: Rhizobiaceae) and “*Candidatus* Phytoplasma spp.” (Bacilli: Acholeplasmatales: Acholeplasmataceae), causing severe problems for agriculture (Jarausch and Jarausch, 2010; Grafton-Cardwell *et al*., 2013; Mora *et al*., 2021). Psyllids depend on vertically transmitted bacterial mutualists to compensate for the nutritional deficiencies in the phloem sap (Profft, 1937; Buchner, 1965; Fukatsu and Nikoh, 1998; Subandiyah *et al*., 2000; Thao, Clark, *et al*., 2000; Thao, Moran, *et al*., 2000; Spaulding and von Dohlen, 2001; Nakabachi *et al*., 2006, 2010; Sloan and Moran, 2012; Nakabachi, Ueoka, *et al*., 2013; Arp *et al*., 2014; Sloan *et al*., 2014; Hall *et al*., 2016; Morrow *et al*., 2017; Nakabachi, Malenovský, *et al*., 2020; Nakabachi, Piel, *et al*., 2020; Kwak *et al*., 2021; Nakabachi *et al*., 2022a, 2022b; Nakabachi and Moran, 2022; Dittmer *et al*., 2023; Maruyama *et al*., 2023; Nakabachi and Suzaki, 2023). They have a specialized organ called the bacteriome (Nakabachi *et al*., 2010; Sloan *et al*., 2014; Nakabachi and Suzaki, 2023), harboring typically two intracellular symbiont species transovarially transmitted through generations. One of them is the primary symbiont “*Candidatus* Carsonella ruddii” (Gammaproteobacteria: Oceanospirillales: Halomonadaceae) (Thao, Moran, *et al*., 2000), providing the host with essential amino acids scarce in the phloem sap (Nakabachi *et al*., 2006; Sloan and Moran, 2012; Nakabachi, Ueoka, *et al*., 2013; Nakabachi, Piel, *et al*., 2020; Dittmer *et al*., 2023), and is thus considered essential for Psylloidea. Molecular phylogenetic analyses demonstrated a single acquisition of a *Carsonella* ancestor by a psyllid common ancestor and its subsequent stable vertical transmission (Thao, Moran, *et al*., 2000; Spaulding and von Dohlen, 2001; Hall *et al*., 2016; Nakabachi, Malenovský, *et al*., 2020; Nakabachi *et al*., 2022b, 2022a; Maruyama *et al*., 2023). Another bacteriome-associated bacterium is classified as a ‘secondary symbiont,’ with wide phylogenetic variations among host lineages, suggesting its repeated replacements during the evolution of Psylloidea (Thao, Clark, *et al*., 2000; Spaulding and von Dohlen, 2001; Sloan and Moran, 2012; Hall *et al*., 2016; Morrow *et al*., 2017; Nakabachi, Malenovský, *et al*., 2020; Nakabachi *et al*., 2022b, 2022a). They are principally believed to have a nutritional basis (Spaulding and von Dohlen, 2001; Sloan and Moran, 2012; Morrow *et al*., 2017; Dittmer *et al*., 2023), with a distinct exception of “*Candidatus* Profftella armatura” (Gammaproteobacteria: Burkholderiales: Oxalobacteraceae) (Nakabachi, Ueoka, *et al*., 2013; Dan *et al*., 2017; Nakabachi, Malenovský, *et al*., 2020; Nakabachi, Piel, *et al*., 2020), whose leading role is considered to be the protection of the holobiont (host-symbiont complex) from natural enemies (Nakabachi, Ueoka, *et al*., 2013; Nakabachi and Fujikami, 2019; Nakabachi and Okamura, 2019; Yamada *et al*., 2019; Nakabachi, Piel, *et al*., 2020; Tanabe *et al*., 2022; Takasu *et al*., 2023, 2024). In addition to these obligate mutualists harbored in the bacteriome, psyllids may be infected with various secondary symbionts of a facultative nature, including *Wolbachia* (Alphaproteobacteria: Rickettsiales: Anaplasmataceae), *Rickettsia* (Alphaproteobacteria: Rickettsiales: Rickettsiaceae), *Diplorickettsia* (Gammaproteobacteria: Diplorickettsiales: Diplorickettsiaceae), and *Spiroplasma* (Bacilli: Entomoplasmatales: Spiroplasmataceae), which may cause systemic infection in the host insects (Spaulding and von Dohlen, 2001; Sloan and Moran, 2012; Arp *et al*., 2014; Chu *et al*., 2016, 2019; Jain *et al*., 2017; Kruse *et al*., 2017; Morrow *et al*., 2017; Nakabachi, Malenovský, *et al*., 2020; Kwak *et al*., 2021; Cooper *et al*., 2022; Nakabachi *et al*., 2022b, 2022a). As with other sap feeders (Nakabachi and Ishikawa, 1997, 1999, 2000, 2001; Nakabachi *et al*., 2003, 2005, 2014; Moran *et al*., 2008; Kikuchi, 2009; Nikoh and Nakabachi, 2009; Shigenobu *et al*., 2010; Tamborindeguy *et al*., 2010; Nakabachi, 2015; Uchi *et al*., 2019), there is growing evidence that interactions among the host and these symbiotic microbes considerably affect psyllid biology and host plant pathology (Nakabachi, Nikoh, *et al*., 2013; Chu *et al*., 2016, 2019; Dan *et al*., 2017; Jain *et al*., 2017; Kruse *et al*., 2017; Nakabachi, Malenovský, *et al*., 2020; Tanabe *et al*., 2022; Killiny, 2022; Nakabachi *et al*., 2022b; Cooper *et al*., 2023). However, little is known about the microbiome of *A. itadori*, which potentially impacts properties as a biocontrol agent, whereas random sequencing of the genomic libraries derived from Kyushu and Hokkaido strains detected sequences similar to those of *Carsonella* and *Sodalis* spp. (Gammaproteobacteria: Enterobacterales: Pectobacteriaceae) (Andersen *et al*., 2016). To address this issue, the present study analyzed the microbiota of an *A. itadori* strain newly collected in Honshu, the central main island of Japan, along with related species belonging to the family Aphalaridae, using amplicon sequencing of 16S rRNA genes. The localization of symbionts in the *A. itadori* body was further analyzed using the fluorescence *in situ* hybridization (FISH) method.

## Materials and Methods

### Insects

For amplicon analyses, adults of four psyllid species belonging to the family Aphalaridae were collected from their host plants at various locations in Japan (Table 1).

**Table 1.**
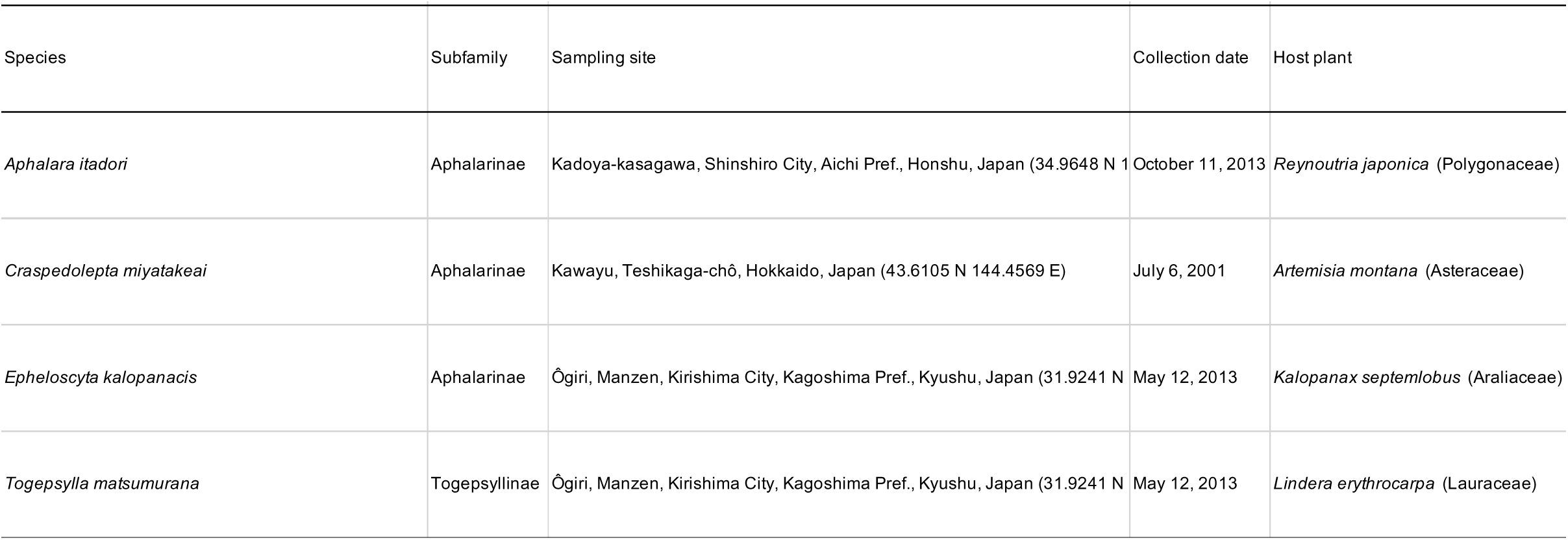
Psyllid species used for the amplicon analysis.

*A. itadori*, *Craspedolepta miyatakeai*, and *Epheloscyta kalopanacis* were selected from Aphalarinae, the most species-rich subfamily of Aphalaridae. In addition, *Togepsylla matsumurana* was chosen from the subfamily Togepsyllinae with distinctive morphological features (Luo *et al*., 2017). The insect samples were stored in acetone (*A. itadori*) or 99.5% ethanol (the other species) at -20°C until DNA extraction. For FISH analysis, *A. itadori* individuals derived from a colony established in the laboratory were used. The colony was originally collected from *R. japonica* in Shinshiro City, Aichi Prefecture, Honshu, Japan (34.9650 N, 137.5729 E: close to the site where specimens used for amplicon analysis were collected) on May 11, 2022. The *A. itadori* population was maintained on *R. japonica* covered with insect-rearing sleeves (L70 cm×W30 cm, Bugdorm Store; Taichung, Taiwan) and kept in incubators set at 20 °C with a photoperiod of 16:8 (L:D) h.

### DNA extraction

DNA was extracted from the whole bodies of pooled individuals of five adult males and five adult females of each psyllid species using the DNeasy Blood & Tissue Kit (Qiagen, Hilden, Germany). The quality of the extracted DNA was assessed using a NanoDrop 2000c spectrophotometer (Thermo Fisher Scientific). The quantity was assessed using a Qubit 2.0 fluorometer with the Qubit dsDNA HS Assay Kit (Thermo Fisher Scientific).

### Construction and sequencing of amplicon libraries

Bacterial populations in psyllids were analyzed using the MiSeq system (Illumina), as described previously (Nakabachi, Malenovský, *et al*., 2020; Nakabachi *et al*., 2022b, 2022a; Maruyama *et al*., 2023). Amplicon polymerase chain reaction (PCR) was performed using DNA extracted from psyllids, KAPA HiFi HotStart ReadyMix (KAPA Biosystems), and the primer set 16S_341Fmod (5′-TCGTCGGCAGCGTCAGATGTGTATAAGAGACAGYYTAMGGRNGGCWGC AG-3′) and 16S_805R (5′-GTCTCGTGGGCTCGGAGATGTGTATAAGAGACAGGACTACHVGGGTATC TAATCC-3′), targeting the V3 and V4 regions of the 16S rRNA gene. Dual indices and Illumina sequencing adapters were attached to the amplicons with index PCR using Nextera XT Index Kit v2 (Illumina). The libraries were combined with the PhiX Control v3 (Illumina), and 250 bp of both ends were sequenced on the MiSeq platform (Illumina) with MiSeq Reagent Kit v2 (500 cycles; Illumina).

### Computational analysis of bacterial populations

The output sequences were processed using the QIIME2 platform (version 2022.11) (Bolyen *et al*., 2019) as described previously (Nakabachi, Malenovský, *et al*., 2020; Nakabachi *et al*., 2022b, 2022a; Maruyama *et al*., 2023). Following primer removal, paired-end reads were denoised and joined, and low-quality or chimeric reads were discarded using the dada2 plugin (Callahan *et al*., 2016). Taxonomic information was assigned to the dereplicated amplicon reads via q2-feature-classifier (Bokulich *et al*., 2018), which had been trained with V3-V4 regions of the 16S rRNA gene (Silva 138 SSURef NR99) (Glöckner *et al*., 2017). Sequence variants (SVs) obtained were manually checked with BLASTN searches against the National Center for Biotechnology Information non-redundant database (Camacho *et al*., 2009). Following the alignment of SVs with related sequences with SINA (version 1.2.11) (Pruesse *et al*., 2012), phylogenetic trees were estimated by the maximum likelihood (ML) method using RAxML (version 8.2.12) (Stamatakis, 2014). The GTR + Γ model was used with no partitioning of the data matrix, with 1,000 bootstrap iterations (options -f a -m GTRGAMMA -# 1000).

### FISH analysis with tissue sections

Adults of *A. itadori* were fixed in Carnoy’s solution (ethanol:chloroform:glacial acetic acid, 6:3:1) at room temperature overnight. After washing with ethanol, the fixed samples were treated with 6% H_2_O_2_ in 80% ethanol until sufficiently decolorized.

Following washing with ethanol, bleached samples were infiltrated and embedded in polyester wax (VWR) and then sliced into serial sections (5 µm thickness) using a rotary microtome RV-240 (Yamato Koki). The sections were mounted on silane-coated Platinum Pro glass slides (Matsunami Glass) and dewaxed in ethanol. The samples were then rehydrated using graded ethanol to phosphate-buffered saline (PBS) series. Probes Car1 (5′-CGCGACATAGCTGGATCAAG-3′) [10] and Sod1 (5′-CGCGGCATGGCTGCATCAGGG-3′) were used to specifically detect 16S rRNA from *Carsonella* and *Sodalis*, respectively. Car1 and Sod1 were 5′-labeled with Alexa Fluor 568 and Alexa Fluor 488, respectively. The tissue sections on glass slides were pre-incubated with hybridization buffer [20 mM Tris-HCl (pH8.0), 0.9M NaCl, 0.01% sodium dodecyl sulfate, 20% formamide], without probes, at room temperature for 1 h. The sections were then incubated at room temperature overnight with a hybridization buffer containing 100 nM of each probe. The samples were washed twice with PBS and mounted in ProLong Gold antifade reagent with DAPI (Thermo Fisher Scientific) using a cover slip. The slides were examined by fluorescence microscopy (BX-53; Olympus) or confocal laser microscopy (A1; Nikon).

### Whole-mount FISH analysis

As mentioned above, fifth instar nymphs of *A. itadori* were fixed and decolorized. After washing with ethanol, samples were hydrated with PBSTx (0.8% NaCl, 0.02% KCl, 0.115% Na_2_HPO_4_, 0.02% KH_2_PO_4_, 0.3% Triton X-100), pre-incubated three times (20 min per incubation) with the hybridization buffer without probes, and then incubated with the hybridization buffer containing 100 nM of each probe at room temperature overnight. Following washing twice with PBSTx, the samples were transferred onto glass slides with spacers and mounted in ProLong Gold antifade reagent with DAPI using a cover slip. The specimens were examined using a Nikon A1 laser scanning confocal microscope, and acquired images were analyzed using NIS-elements AR Analysis 4.10 software (Nikon).

### Data availability

The nucleotide sequence data are available in the DDBJ/EMBL/GenBank databases under the accession numbers DRR437490–DRR437493 (MiSeq output) and TAAR01000001–TAAR01000011 (dereplicated SVs).

## Results

### All four Aphalaridae species have Carsonella

MiSeq sequencing of the amplicon libraries yielded 41,992–83,361 pairs of forward and reverse reads for the four psyllid species. Denoising and joining the paired-end reads with discarding low-quality or chimeric reads resulted in 36,180–78,303 nonchimeric high-quality reads (Table S1). These reads were dereplicated into 227 independent SVs, among which 11 SVs accounted for >1% of the total reads (Table S2). This study focused on these 11 main SVs unless otherwise stated because filtering with a threshold of 1% is demonstrated to be among the most effective and accurate methods to remove potential contaminants (Karstens *et al*., 2019). SVs with a relative abundance of <1% are collectively classified as ‘others’ in Fig. 1, which account for 0.9% (*Togepsylla matsumurana*)–15.6% (*Craspedolepta miyatakeai*) of the total reads in each psyllid species (Table S2). Taxonomic classification by QIIME2 (Table S2) followed by independent BLAST searches and phylogenetic analyses showed that all psyllid species analyzed in this study harbor distinct lineages of *Carsonella* (Gammaproteobacteria: Oceanospirillales: Halomonadaceae, Figs. 1 and 2). The ML tree placed these *Carsonella* sequences at positions that are consistent with the host psyllid phylogeny inferred by mitochondrial and nuclear gene analyses with the aid of morphological analyses (Percy *et al*., 2018; Cho *et al*., 2019; Burckhardt *et al*., 2021), providing further support for the hypothesis that a *Carsonella* ancestor was acquired by a psyllid common ancestor followed by stable vertical transmission (Fig. 2).

**Fig. 1.**
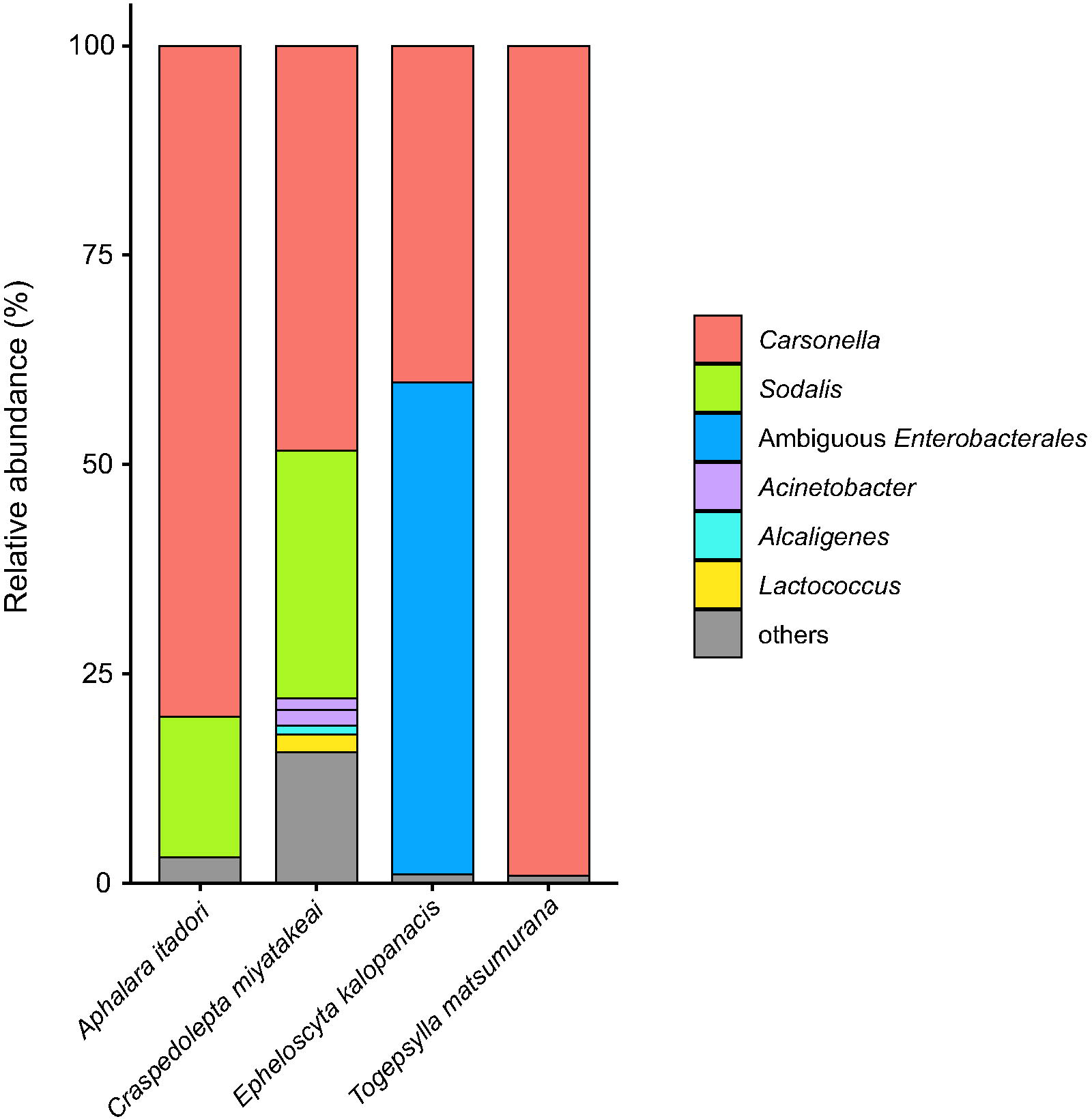
Composition of bacterial populations in psyllids of the family Aphalaridae. Relative abundances of Illumina reads belonging to assigned bacterial taxa are shown.

**Fig. 2.**
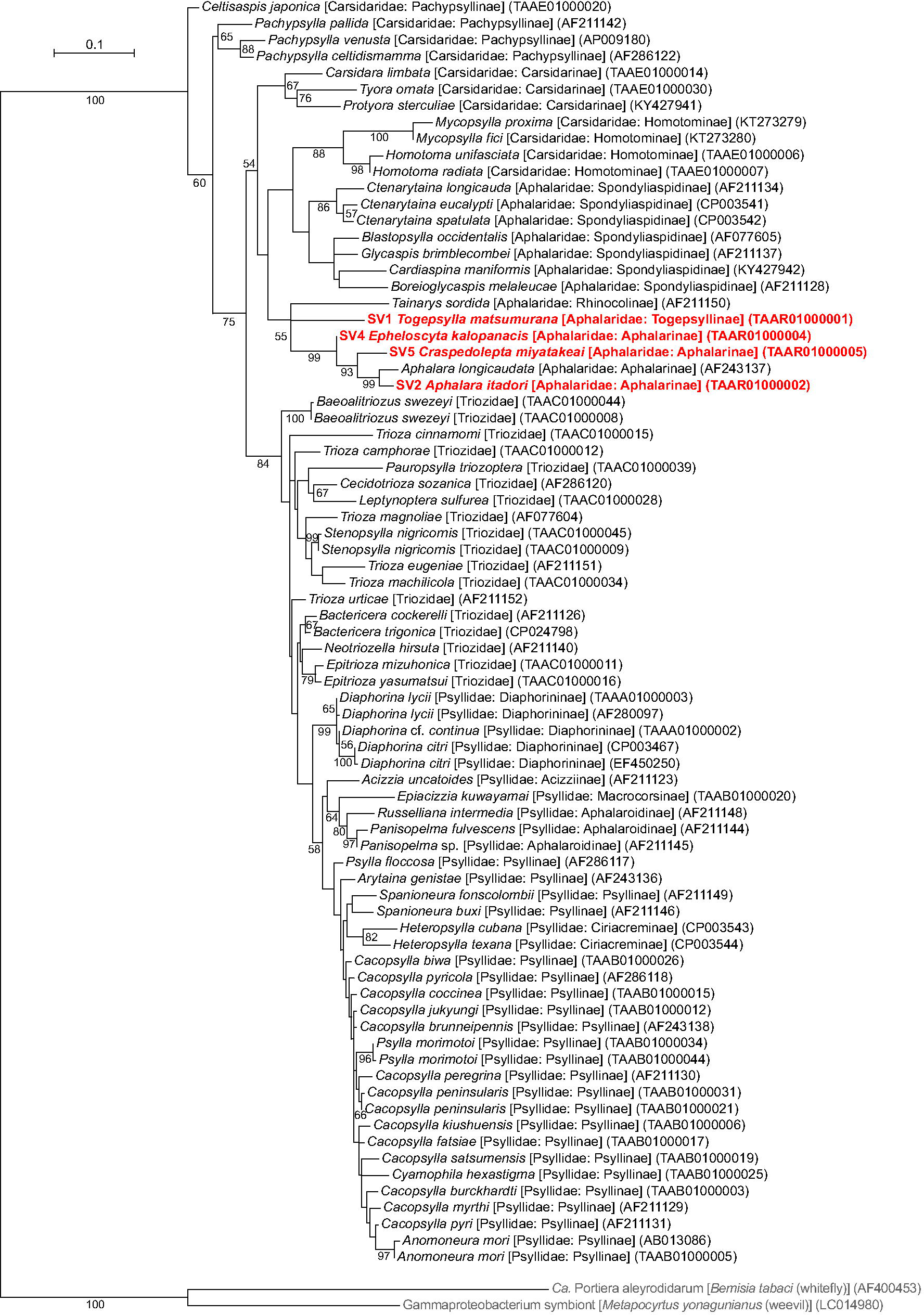
Maximum likelihood phylogram of *Carsonella*. A total of 427 unambiguously aligned nucleotide sites of 16S rRNA genes were subjected to the analysis. On each branch, bootstrap support values of >50% are shown. Designations other than those for outgroups refer to psyllid hosts. Families and subfamilies (if applicable) of host psyllids are shown in brackets. Sequences from this study are shown in bold red. DDBJ/EMBL/GenBank accession numbers for sequences are provided in parentheses. The bar represents nucleotide substitutions per position. The outgroups were “*Ca*. Portiera aleyrodidarum;” the primary symbiont of the whitefly *Bemisia tabaci* (Hemiptera: Sternorrhyncha: Aleyrodoidea), and a gammaproteobacterium symbiont of the weevil *Metapocyrtus yonagunianus* (Coleoptera: Curculionidae).

### A. itadori *has a* Sodalis *symbiont*

The analysis demonstrated that *A. itadori* harbors *Sodalis* sp. (Gammaproteobacteria: Enterobacterales: Pectobacteriaceae) as a secondary symbiont (Figs. 1 and 3, Table S2). Namely, QIIME2 assigned SV2 and SV7, which accounted for 80.1% and 16.8%, respectively, of *A*. *itadori* reads, to *Carsonella* and *Sodalis* (Fig. 1, Table S2). These were the only SVs accounting for >1% of the total reads in *A*. *itadori*, suggesting that *Sodalis* is essentially the only secondary symbiont in the analyzed strain of *A*. *itadori.* Moreover, no apparent plant pathogens, including “*Candidatus* Liberibacter spp.” and “*Candidatus* Phytoplasma spp.”, or facultative symbionts with the parasitic nature, including *Wolbachia*, *Rickettsia*, *Diplorickettsia*, and *Spiroplasma*, were found in the strain even when the search was extended to include SVs with a relative abundance of <1% (Table S2). The top BLAST hit of SV7 was the sequence of *Sodalis* endosymbiont (AB915777) of the stinkbug *Nezara antennata* (Hemiptera: Heteroptera: Pentatomidae). SV7 was 98.1% identical to that of “*Candidatus* Sodalis pierantonius” (AF548137), the primary symbiont of the rice weevil *Sitophilus oryzae* (Coleoptera: Curculionidae) (Dale *et al*., 2003), and 97.9% identical to the sequence of the type species *Sodalis glossinidius* (AP008232), a secondary symbiont of the tsetse fly *Glossina morsitans* (Diptera: Glossinidae) (Dale and Maudlin, 1999). SV7 formed a clade with *Sodalis* symbionts found in various insects, including those of other psyllid species, *Homotoma unifasciata* (Carsidaridae: Homotominae; TAAE01000023, TAAE01000026, and TAAE01000031, 95.8% – 96.7% identical to SV7) (Maruyama *et al*., 2023), *Cacopsylla kiushuensis* (Psyllidae: Psyllinae; TAAB01000030, 96.5% identical to SV7), *Cacopsylla burckhardti* (Psyllidae: Psyllinae; TAAB01000016, 96.7% identical to SV7) (Nakabachi *et al*., 2022b), and *C. miyatakeai* (Aphalaridae: Aphalarinae; TAAR01000006=SV6, 96.7% identical to SV7, see below) (Fig. 3). Although the clade consisting of *Sodalis* symbionts was only poorly supported (bootstrap: < 50%) (Fig. 3), SV7 was tentatively named “*Sodalis* endosymbiont” (Figs. 1 and 3) because its similarities to the type species was above the generally used arbitrary genus threshold of 94.5–95% (Yarza *et al*., 2014; Barco *et al*., 2020).

**Fig. 3.**
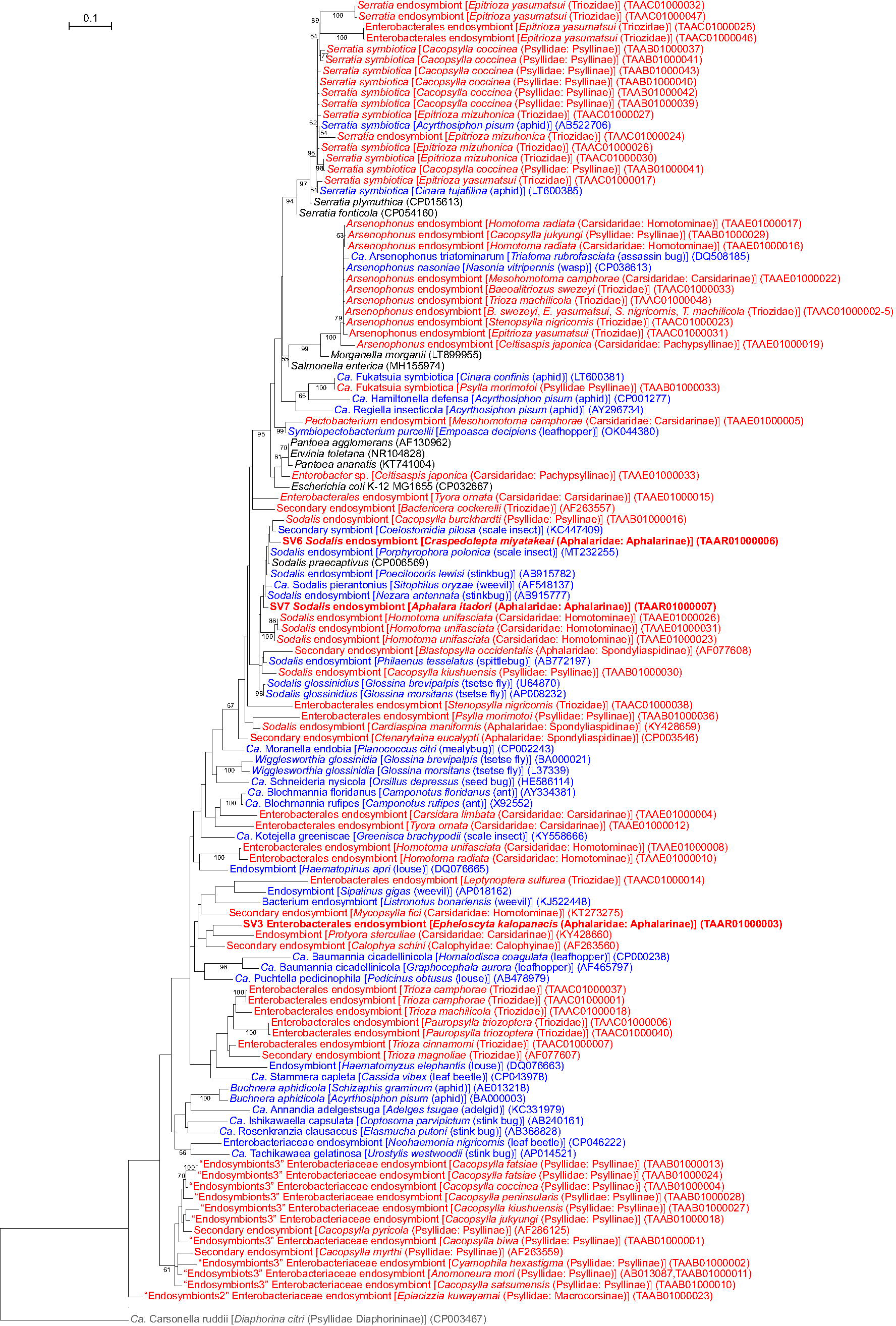
Maximum likelihood phylogram of *Enterobacterales*. A total of 427 unambiguously aligned nucleotide sites of 16S rRNA genes were subjected to the analysis. On each branch, bootstrap support values of > 50% are shown. The scale bar indicates substitutions per site. For symbiotic bacteria, host organisms are shown in brackets. Symbionts of animals other than psyllids are shown in blue, whereas symbionts of psyllids are shown in red. Sequences from this study are shown in bold. DDBJ/EMBL/GenBank accession numbers are provided in parentheses. *Carsonella* was used as an outgroup.

### C. miyatakeai *has* Sodalis *not sister to the* A. itadori *counterpart*

QIIME2 assigned SV5 and SV6, which accounted for 48.4% and 29.6% of *C*. *miyatakeai* reads to *Carsonella* and *Sodalis*, respectively (Fig. 1, Table S2). SV6 was 96.3% identical to the sequence of the type species *S. glossinidius* (AP008232) and was thus also tentatively named “*Sodalis* endosymbiont.” In the ML tree, SV6 did not form an exclusive clade with SV7, corresponding to the *Sodalis* endosymbiont of *A*. *itadori*. Instead, these SVs formed a large clade with a low bootstrap support value (< 50%), interrupted by numerous sequences of other *Sodalis* lineages derived from various insect hosts and even *Sodalis praecaptivus*, which was isolated from a human wound (Chari *et al*., 2015) (Fig. 3). This branching pattern indicates that the *Sodalis* symbionts of *A*. *itadori* and *C*. *miyatakeai* are not sister lineages, suggesting that each host lineage independently acquired them.

SV8, which accounted for 2.2% of *C*. *miyatakeai* reads, was 100% identical to the sequence of *Lactococcus lactis* (Bacilli: Lactobacillales: Streptococcaceae; e.g. CP069223). SV9 and SV10, accounting for 1.9% and 1.4% of *C*. *miyatakeai* reads, were 100% identical to sequences of *Acinetobacter* spp. (Gammaproteobacteria: Pseudomonadales: Moraxellaceae). SV11, representing 1.1% of *C*. *miyatakeai* reads, was 100% identical to sequences of *Alcaligenes* spp. (Gammaproteobacteria: Burkholderiales: Alcaligenaceae; e.g. MT572474) (Figs. 1 and 3, Table S2). These low relative abundances and the perfect identity with free-living bacteria implied that the corresponding bacteria were transient associates, not stable symbionts.

### E. kalopanacis has a non-Sodalis secondary symbiont

QIIME2 assigned SV3 and SV4, which accounted for 58.8% and 40.2% of *E. kalopanacis* reads, to an ambiguous *Enterobacterales* symbiont (Gammaproteobacteria: Enterobacterales: Morganellaceae) and *Carsonella*, respectively (Figs. 1 and 3, Table S2). The top BLAST hit of SV3 was the sequence of “*Candidatus* Schneideria nysicola” (Gammaproteobacteria: Enterobacterales: Morganellaceae) (HE586114, 92.5% identical to SV3), an endosymbiont of the seed bug *Orsillus depressus* (Hemiptera: Heteroptera: Lygaeidae) (Kuechler *et al*., 2012), which was followed by sequences of symbionts derived from other psyllid species, *Calophya schini* (Calophyidae: Calophyinae, AF263560, 92.3% identical to SV3) (Thao, Clark, *et al*., 2000) and *Protyora sterculiae* (Carsidaridae: Carsidarinae, KY428660, 91.9% identical to SV3) (Morrow *et al*., 2017). Although SV3 and the latter two sequences formed a poorly supported clade (bootstrap < 50%), their overall phylogenetic position was ambiguous within the order of Enterobacterales (Fig. 3).

### No secondary symbionts were found in T. matsumurana

QIIME2 assigned SV1, which accounted for 99.1% of *T. matsumurana* reads, to *Carsonella*. No other sequence with a relative abundance of >1% was detected in *T. matsumurana* (Figs. 1 and 2, Table S2).

### A. itadori harbors Sodalis in the bacteriome

FISH analyses demonstrated that the *Sodalis* symbiont identified in the amplicon analysis is harbored along with *Carsonella* in the *A. itadori* bacteriome (Figs. 4 and S1), suggesting that not only *Carsonella* but also *Sodalis* is an evolutionarily stable mutualist for *A. itadori*. The whole-mount analysis using the fifth instar nymphs showed that *Sodalis* cells are localized in the central area of the bacteriome surrounded by bacteriocytes harboring *Carsonella* (Figs. 4A and S1). Although this localization of the secondary symbiont and the overall shape of the bilobed bacteriome were similar to those of other psyllid species reported previously (Profft, 1937; Buchner, 1965; Nakabachi, Ueoka, *et al*., 2013; Dan *et al*., 2017), the central area for harboring the secondary symbiont appeared much smaller. This is especially the case when compared with the bacteriome of the same developmental stage, fifth instar nymph, of the Asian citrus psyllid *Diaphorina citri* (Psyllidae: Diaphorininae), where the bacteriome mainly consisted of the central area, only peripherally margined by the bacteriocytes for *Carsonella* (Dan *et al*., 2017). FISH analysis using sectioned tissues of *A. itadori* adults also clearly showed that *Sodalis* is localized and densely packed in the central area of the bacteriome (Fig. 4B). The analysis further suggested that the central area consists of multiple uninucleate bacteriocytes with a nucleus larger than those of the bacteriocytes for *Carsonella*.

**Fig. 4.**
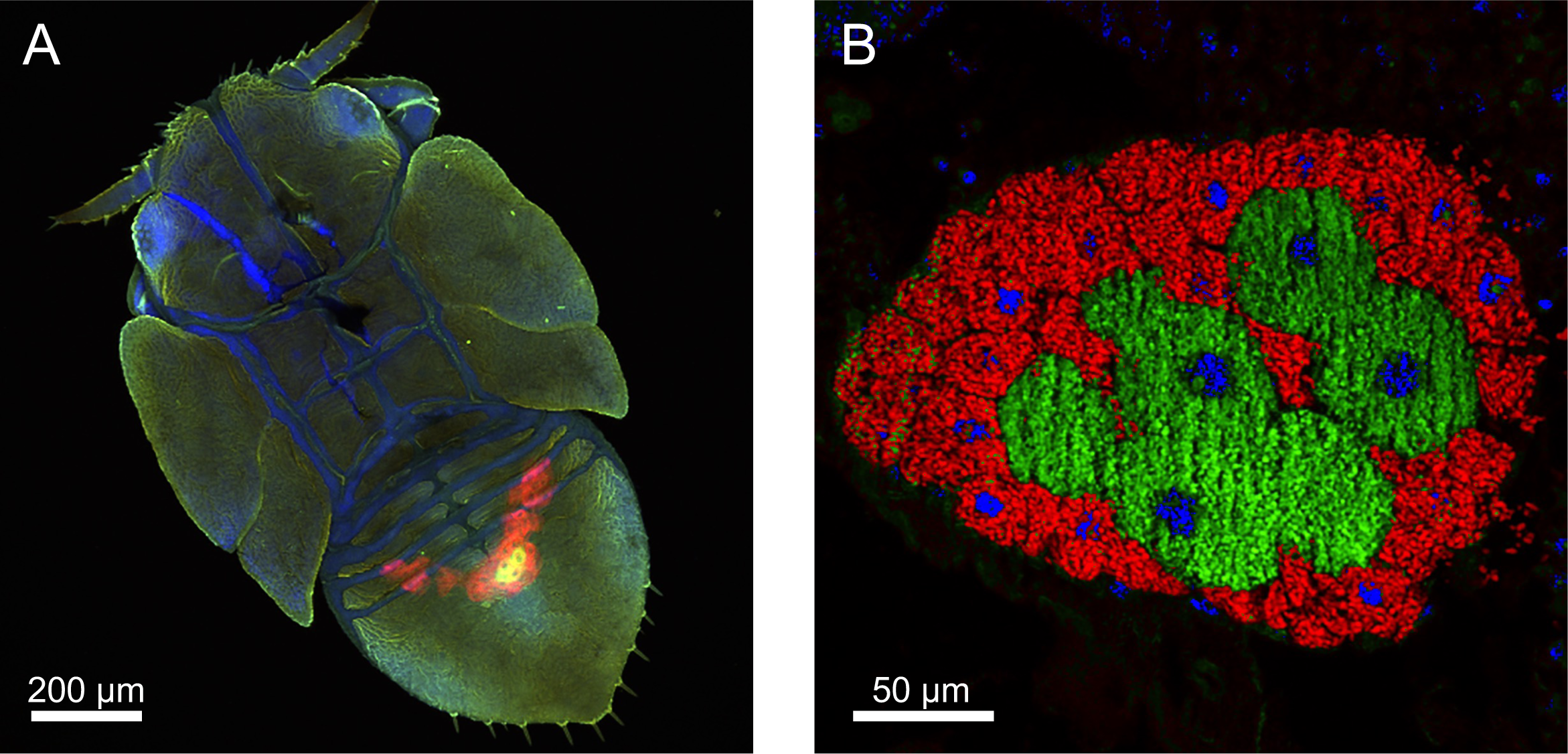
FISH images of bacteriomes of *A. itadori*. Red (Alexa Fluor 568), green (Alexa Fluor 488), and blue (DAPI) signals indicate *Carsonella*, *Sodalis*, and the host nuclei, respectively. (A) Whole-mount FISH image [maximum intensity projection (MIP)] of the dorsal view of a fifth instar nymph showing the bacteriome in the abdomen. (B) FISH image of a cross-section of the abdomen of an adult female. *Carsonella* is harbored within the bacteriocytes on the surface of the bacteriome, while *Sodalis* is encased in the other type of bacteriocytes with large nuclei, located at the center of the bacteriome.

## Discussion

This study demonstrated that the *A. itadori* strain newly collected on the island of Honshu has a dual symbiotic system involving *Carsonella* and *Sodalis* (Figs. 1–4, Table S2). FISH analysis of *A. itadori* further demonstrated that both *Carsonella* and *Sodalis* are harbored in the bacteriome (Figs. 4 and S1), suggesting that these bacteria are evolutionarily stable mutualists for *A. itadori*, which is consistent with the previous report that random sequencing of the genomic libraries derived from Kyushu and Hokkaido strains detected sequences similar to those of *Carsonella* and *Sodalis* (Andersen *et al*., 2016). While *Carsonella* was housed in uninucleate bacteriocytes peripherally located in the bacteriome, the *Sodalis* symbiont was shown to be harbored in the central area of the bacteriome (Figs. 4 and S1). It has been believed that the bacteriome of psyllids generally consists of a central syncytium to harbor a secondary symbiont, uninucleate bacteriocytes to harbor *Carsonella*, and an envelope encasing the whole, although the distribution of the syncytium and bacteriocytes varies slightly depending on the psyllid taxa (Profft, 1937; Buchner, 1965; Chang and Musgrave, 1969; Waku and Endo, 1987; Fukatsu and Nikoh, 1998; Subandiyah *et al*., 2000; Nakabachi, Ueoka, *et al*., 2013; Dan *et al*., 2017; Dittmer *et al*., 2023; Nakabachi and Suzaki, 2023). However, our observation strongly suggested that the central area of the *A*. *itadori* bacteriome is not a syncytium but consists of uninucleate bacteriocytes with features distinct from those of bacteriocytes for harboring *Carsonella*. The bacteriocytes for *Sodalis* appeared to be larger in size, having a nucleus also larger than those of bacteriocytes for *Carsonella* (Fig. 4B), suggesting that the former has higher ploidy than the latter (Nakabachi *et al*., 2010). This larger nuclear size also contrasts with the cases of the previously reported bacteriomes of other psyllid species, in which the syncytium has smaller nuclei than those of the bacteriocytes for *Carsonella* (Profft, 1937; Buchner, 1965; Nakabachi, Ueoka, *et al*., 2013; Dan *et al*., 2017; Dittmer *et al*., 2023; Nakabachi and Suzaki, 2023). Studies on the embryonic development of several psyllid lineages showed that the syncytium of the bacteriome is formed through fusion of provisional uninucleate bacteriocytes that already harbor secondary symbionts (Profft, 1937; Buchner, 1965; Dan *et al*., 2017). Therefore, during the development of *A. itadori*, the immature uninucleate bacteriocytes harboring *Sodalis* may exceptionally continue to grow increasing their ploidy, without fusing to form the syncytium. Further studies are required to assess this hypothesis.

*Sodalis* symbionts have been found in various insect taxa, including tsetse flies (Dale and Maudlin, 1999), weevils (Toju *et al*., 2013; Oakeson *et al*., 2014), lice (Psocodea: Anoplura) (Fukatsu *et al*., 2007), louse flies (Diptera: Hippoboscoidea) (Nováková and Hypša, 2007), stinkbugs (Hemiptera: Heteroptera) (Kaiwa *et al*., 2010), spittlebugs (Hemiptera: Auchenorrhyncha: Cercopoidea) (Koga *et al*., 2013; Koga and Moran, 2014), scale insects (Hemiptera: Sternorrhyncha: Coccoidea) (Dhami *et al*., 2013; Husnik and McCutcheon, 2016), and psyllids (Arp *et al*., 2014; Hall *et al*., 2016; Morrow *et al*., 2017; Nakabachi *et al*., 2022b; Maruyama *et al*., 2023). Although their symbiotic types can vary across the mutualism-parasitism continuum, the main role of *Sodalis* symbionts appears to be provisioning of nutrients in many insect hosts (McCutcheon *et al*., 2019). Moreover, microbiomic, genomic, and physiological studies suggested that *Sodalis* symbionts have replaced more ancient predecessor symbionts in weevils (Oakeson *et al*., 2014) and spittlebugs (Koga and Moran, 2014), playing the role in supplementing nutrients formerly supplied by the predecessor symbionts (Koga and Moran, 2014; Oakeson *et al*., 2014). Because similar symbiont replacements by *Sodalis* were also suggested in psyllid lineages (Nakabachi *et al*., 2022b), their functional roles in Psylloidea, especially *A. itadori*, are of interest.

Although no direct evidence exists that *A. itadori* harms plants other than *Reynoutria* spp. (Shaw *et al*., 2009), adult psyllids also feed on plants other than true hosts on which the immature to adult life cycle is completed (Burckhardt *et al*., 2014; Cooper *et al*., 2019), posing a potential risk of spreading plant pathogens. Moreover, evidence is accumulating that facultative symbionts manipulating host reproduction are frequently horizontally transmitted among various arthropod hosts (Porter and Sullivan, 2023). In this context, the *A. itadori* strain analyzed in this study was demonstrated to have no apparent plant pathogens, including “*Candidatus* Liberibacter spp.” and “*Candidatus* Phytoplasma spp.,” or facultative symbionts with a parasitic nature, including *Wolbachia*, *Rickettsia*, *Diplorickettsia*, and *Spiroplasma* (Table S2), indicating that this strain is suitable as a biocontrol agent imposing a minimum risk on the ecosystem.

This study identified various bacterial populations in psyllid species of the family Aphalaridae. Although a *Sodalis* symbiont was also identified in *C*. *miyatakeai* (Fig. 1, Table S2), phylogenetic analysis indicated that these *Sodalis* symbionts were independently acquired by each psyllid lineage (Fig 3), providing further support for the hypothesis that bacteriome-associated secondary symbionts have been repeating infection and replacement during the evolution of Psylloidea.

## Conclusions

This study revealed that the *A. itadori* strain newly collected in Honshu has a dual symbiotic system involving *Carsonella* and *Sodalis.* While *Carsonella* was harbored in the uninucleate bacteriocytes peripherally located in the bacteriome, *Sodalis* was housed in large uninucleate bacteriocytes with large nuclei located centrally in the bacteriome. No apparent plant pathogens or manipulators of arthropod reproduction were detected in the analyzed strain, indicating that the strain is suitable as a biocontrol agent, posing a minimum risk to the ecosystem.

## Acknowledgements

This work was supported by the Japan Society for the Promotion of Science (https://www.jsps.go.jp) KAKENHI (grant numbers 21687020, 26292174 and 20H02998 to AN, and 25850035 to HI). The funders had no role in the study design, data collection and analysis, decision to publish, or manuscript preparation. The DNA sequencing facility was supported by the Department of Applied Chemistry and Life Science and the University-Community Partnership Promotion Center at the Toyohashi University of Technology.

## Conflict of interest

The authors declare no conflict of interest.

---

**Fig. S1** (mp4, 2.5 MB) Three-dimensional reconstruction of the *A. itadori* body corresponding to the FISH image shown in Fig. 4A.

**Table S1** (xlsx, 11 KB) Processing of sequence reads with QIIME2.

**Table S2** (xlsx, 59 KB) Sequence variants detected in the present study.

## Notes

### Competing Interest Statement

The authors have declared no competing interest.

